# A deep mutational scanning platform to characterize the fitness landscape of anti-CRISPR proteins

**DOI:** 10.1101/2021.08.21.457204

**Authors:** Tobias Stadelmann, Daniel Heid, Michael Jendrusch, Jan Mathony, Stéphane Rosset, Bruno E. Correia, Dominik Niopek

## Abstract

Deep mutational scanning is a powerful method to explore the mutational fitness landscape of proteins. Its adaptation to anti-CRISPR proteins, which are natural CRISPR-Cas inhibitors and key players in the co-evolution of microbes and phages, would facilitate their in-depth characterization and optimization. Here, we developed a robust anti-CRISPR deep mutational scanning pipeline in *Escherichia coli* combining synthetic gene circuits based on CRISPR interference with flow cytometry-coupled sequencing and mathematical modeling. Using this pipeline, we created and characterized comprehensive single point mutation libraries for AcrIIA4 and AcrIIA5, two potent inhibitors of *Streptococcus pyogenes* Cas9. The resulting mutational fitness landscapes revealed that both Acrs possess a considerable mutational tolerance as well as an intrinsic redundancy with respect to Cas9 inhibitory features, suggesting evolutionary pressure towards high plasticity and robustness. Finally, to demonstrate that our pipeline can inform the optimization and fine-tuning of Acrs for genome editing applications, we cross-validated a subset of AcrIIA4 mutants via gene editing assays in mammalian cells and *in vitro* affinity measurements. Together, our work establishes deep mutational scanning as powerful method for anti-CRISPR protein characterization and optimization.

## INTRODUCTION

Deep mutational scanning (DMS) explores next-generation sequencing (NGS) technologies to measure the effect of mutations on proteins in high throughput. It facilitates the creation of comprehensive mutational fitness landscapes, which allow deducing various protein functional and structural properties and can inform protein optimization and engineering (1). In general, DMS requires (i) a robust pipeline for generating a mutant library, (ii) a coupled genotype-phenotype platform and corresponding assay to separate active from inactive protein variants and (iii) an NGS-based readout to assess the enrichment of variants in the active and/or inactive variant pools.

In the past, DMS has been applied, for instance, to characterize human disease mutations (2), dissect protein-coding regions in the adeno-associated virus genome (3) and investigate the mutational fitness landscape of CRISPR-Cas9 (4,5). Moreover, comprehensive DMS datasets comprising protein single and double mutants were used for detailed analysis of intramolecular interactions, based on which the three-dimensional protein structure could be inferred (6). Very recently, DMS was also used to analyze the impact of mutations in the SARS-CoV-2 receptor binding domain on the binding of ACE2, a central host factor required for virus entrance (7). Overall, these studies showcase the power of mutational scanning for protein functional dissection and structure-mechanistic analysis.

Anti-CRISPR (Acr) proteins are natural inhibitors of CRISPR-Cas systems (8,9) and play an important role in the co-evolution of phages and their bacterial hosts (10). Hundreds of experimentally validated anti-CRISPR proteins have been described and many thousands have been additionally predicted (11). Together, these Acrs target all common Cas9 orthologues (12-21), as well as Cas12a, a type V CRISPR effector (22,23), and Cas13, a type VI CRISPR effector (24,25).

Since Acrs are central players in the evolutionary battle between phages and their microbial hosts (10,26-28), studying their CRISPR inhibitory mechanisms, structural diversity as well as origin is of key interest to the CRISPR and microbiology research communities. With respect to their evolution, it is hypothesized that different Acr families emerged independently and from unrelated ancestor genes. One could speculate, that Acrs should be under evolutionary pressure not only to robustly inhibit the CRISPR adaptive immune response within the host cell, but likewise to maintain the capacity for rapid adaptation to evolutionary alterations occurring within its cognate Cas effector(s) as well as changes in external factors (e.g. temperature, pH, cell context).

Apart from their great relevance to micro- and phage biology, their small size and - in many cases - robust CRISPR inhibitory function across a broad range of cell types render Acrs highly attractive tools for CRISPR-Cas control in gene editing applications. Acrs can, for instance, be employed as parts in CRISPR-Cas-based gene circuits (29), were shown to enable reduction of off-target editing via temporally confined and fine-tuned Cas9 activity (30,31), and have been coupled to exogenous or endogenous stimuli such as light (32,33) or microRNAs (34-36) for inducible and cell-type restrictive genome editing, respectively.

Adapting DMS to Acrs could expand both, our knowledge on Acr functional mechanism and evolution, as well as their utility for CRISPR-Cas regulation in context of genome editing applications. Comprehensive Acr mutational fitness landscapes could, for instance, provide important insights into the mutational tolerance of Acrs and hence into their evolutionary design space, and also inform mechanistic studies of newly identified Acrs. Moreover, Acr mutational fitness landscapes could guide the engineering of customized Acrs with altered or improved inhibition potency or specificity (34). Finally, the ability to selectively alter regions of anti-CRISPR polypeptides by mutagenesis without perturbing their actual function could be important for their *in vivo* application, i.e. to overcome potential issues of immunogenicity similar to previous effort for CRISPR nucleases (37-39).

Here, we present a robust experimental and computational pipeline to examine the mutational fitness landscape of anti-CRISPR proteins via deep mutational scanning. By applying our pipeline to AcrIIA4 (19) and AcrIIA5 (14), both targeting *Streptococcus pyogenes* (*Spy*)Cas9, we created comprehensive datasets on the impact of individual amino acid exchanges on CRISPR-Cas9 inhibition as well as the overall Acr mutational tolerance. We then aligned our DMS data to known sequence and structural features, thereby further characterizing the observed mutational tolerance. Finally, we selected several mutations that improve, preserve or modulate Acr function and cross-validated their activity in mammalian cell gene editing assays as well as *in vitro* affinity measurements. Our work establishes mutational scanning as a powerful strategy for Acr characterization and engineering.

## MATERIAL AND METHODS

### Molecular cloning

All plasmids used in this study are listed in Supplementary Table 1, sgRNA target sequences are shown in Supplementary Table 2. Sequences and maps for the CRISRPi selection vectors are available as Supplementary data 1 (Genbank files). Oligonucleotides and synthetic double-stranded DNA fragments were obtained from Integrated DNA Technologies. PCRs were performed with Q5 Hot Start High Fidelity Polymerase (New England Biolabs) or Invitrogen Platinum Superfi II DNA Polymerase (Thermo Fisher Scientific). Expression vectors were created using classical restriction enzyme cloning or Golden-Gate assembly (40). Restriction enzymes were obtained from New England Biolabs, T4 DNA ligase was obtained from Jena Biosciences. Agarose gel electrophoresis was used to analyze PCR and restriction products. Bands of the expected size were cut out and DNA was extracted with a ZymoClean Gel DNA recovery kit (Zymo Research). For all cloning steps, chemically competent *E. coli* K12 DH5α cells (New England Biolabs) were used. Antibiotics were used at the following concentrations: carbenicillin, 50 µg mL^−1^; chloramphenicol, 25 µg mL^−1^; kanamycin, 50 µg mL^−1^. Plasmid DNA was purified with the ZR Plasmid Miniprep or ZymoPure II Midiprep kit (both Zymo Research). The integrity of all plasmids was verified by Sanger Sequencing (GENEWIZ Europe).

AcrIIA4 (*Listeria monocytogenes)* (19) and AcrIIA5 (Phage D4276) (14) coding sequences were codon optimized for expression in *E. coli*, obtained as double stranded gene fragments and cloned into pBAD24 (pBAD24-sfGFPx1 was a gift from Sankar Adhya & Francisco Malagon, Addgene plasmid #51558) (41) via unique EcoRI and HindIII restriction sites, resulting in the plasmids pBAD24-AcrIIA4 and pBAD24-AcrIIA5.

To construct plasmids co-expressing d*Spy*Cas9 and sgRNA scaffolds, a second multiple cloning site was introduced into pBbA5 (pBbA5c-RFP was a gift from Jay Keasling, Addgene plasmid #35281) (42) by PCR, resulting in pBbA5C_sgMCS. A fragment encoding an *E. coli* codon optimized d*Spy*Cas9 was PCR amplified from vector pdCas9 (gift from Luciano Marraffini, Addgene plasmid #46569) (43) and cloned into pBbA5C_sgMCS using EcoRI and BamHI restriction sites, thereby generating pBbA5C_sgMCS_d*Spy*Cas9. A sgRNA expression cassette with an RFP reporter-targeting spacer sequence was ordered as overlapping DNA oligonucleotides, extended by PCR and cloned into pBbA5c-spCas9 using NotI and SalI restriction sites, resulting in pBbA5C_sgMCS_d*Spy*Cas9_RFP_guide. To generate the RFP reporter plasmid, a bacterial codon optimized RFP encoding sequence was PCR amplified from pBbA5c-RFP; the Anderson Promoter J23102 (http://parts.igem.org/Promoters/Catalog/Anderson) was included in the 5’-extension of the forward primer. A sequence encoding a SsrA degradation tag (44) (AANDENYADAS; corresponds to part BBa_M0052 in iGEM parts registry - parts.igem.org) was included as 5’- extension into the reverse primer to reduce the half-live of RFP in bacterial cells. The resulting PCR fragments were cloned into pJUMP27 (pJUMP27-1AsfGFP was a gift from Chris French, Addgene plasmid #126974) (45) via XbaI and PstI restriction sites, thereby resulting in pJUMP27_J23102_RFP_M0052.

The single codon mutational libraries of AcrIIA4 and AcrIIA5 were generated by back-to-back PCR on pBAD24-AcrIIA4 and -AcrIIA5 with forward primers containing NNB overhangs as previously described (46). PCRs were performed for each position individually to avoid PCR bias, followed by gel extraction. The purified fragments were treated with KLD enzyme mix (New England Biolabs) and transformed into *E. coli* DH5a chemically competent cells. Cells were grown in LB supplemented with carbenicillin. Note that sub-libraries corresponding to a single codon were grown individually and to stationary phase in a 96 deep-well plate. Cultures were then combined at an equal volume and further grown in 50 ml LB carbenicillin until saturation, followed by extraction of plasmid DNA.

Single mutant Acr variants for bacterial validation experiments were created via back-to-back PCR on template vectors pBAD24-AcrIIA4 and -5 and by incorporating the mutations into the 5’-extension of one primer, followed by KLD treatment.

The vectors co-encoding firefly luciferase, a firefly luciferase gene targeting sgRNA and *Renilla* luciferase (for normalization purposes) used for the dual luciferase assay experiments in mammalian cells were previously reported by us (32). Vector pCMV-AcrIIA4 was previously reported by us (32). Single codon substitutions were introduced into the Acr expression constructs via back-to-back PCR by incorporating the mutations into the 5’- extension of one primer, followed by KLD treatment.

### Transformation of *E. coli* with Acr libraries

For library generation, chemically competent bacterial cells were first co-transformed with the d*Spy*Cas9 plasmid and the RFP reporter plasmid and selected on LB containing kanamycin and chloramphenicol. An overnight culture of a single bacterial colony was inoculated in Super Optimal Broth (SOB) medium supplemented with kanamycin and chloramphenicol, grown to an optical density (OD)_600_ of 0.5 and chemical competent cells were prepared according to the Inoue transformation protocol (47). AcrIIA4 and AcrIIA5 libraries were transformed into chemical competent cells by heat-shock at 42°C for 1 min. 0.1% of the total transformation volume was plated on LB Agar plates containing carbenicillin, chloramphenicol and kanamycin to estimate the transformation efficiency and corresponding library complexity (Supplementary Tables 3 and 4). The remaining cells were grown in liquid LB medium containing carbenicillin, chloramphenicol and kanamycin until stationary phase, followed by cryopreservation in aliquots.

### Fluorescence-activated cell sorting

Cryopreserved cells were thawed and grown in LB medium containing carbenicillin, chloramphenicol, kanamycin until stationary phase, followed by induction with 1 mM IPTG and 4 mM arabinose at a starting OD_600_ of 0.05. Following a 12-hour incubation period, cells were collected by centrifugation at 5000×g for 10 minutes, the supernatant was removed, and cells were resuspended in 10 volumes 1×PBS buffer.

Fluorescence-activated cell sorting (FACS) experiments were performed on a BD FACSAria™ *Fusion* flow cytometer at the ZMBH flow cytometry core facility (Heidelberg University). *E. coli c*ells were first gated using the forward-scatter area (FSC-A) and site-scatter area (SSC-A). Bacterial cells were then sorted into eight fractions according to their RFP intensity using 150,000 total cells per fraction. Collected cells were either frozen for DNA extraction followed by deep amplicon sequencing or recovered by growing the cells in LB medium containing the necessary antibiotics for subsequent characterization of fluorescence enrichments. In the latter case, cells were washed in PBS twice and grown in LB medium containing carbenicillin, chloramphenicol, kanamycin overnight. The cultures form the individual fractions were then induced with 1 mM IPTG and 4 mM arabinose at a starting OD_600_ of 0.05 and analyzed on a BD FACSAria™ *Fusion* flow cytometer 12 hours later as described above.

### Amplicon Deep Sequencing

FACS-sorted *E. coli* cell fractions were lysed and DNA extracted for subsequent 1^st^ stage PCR amplification of AcrIIA4 or AcrIIA5 genes with primers containing Nextera XT index overhangs. PCR amplicons were purified by gel extraction and the concentrations of each sample was adjusted to 25 ng/µl for the 2nd stage barcoding PCR using TG Nextera XT Index Kit v2 Set A (Illumina) according to the manufacturer’s protocol. Libraries were sequenced at the EMBL Heidelberg GeneCore facility on the Illumina MiSeq system using 2×250 paired-end sequencing reagents (MiSeq Reagent Kit v2, 500-cycles).

### NGS data analysis

Paired-end reads for each sorted fraction were assembled by their overlap and filtered to remove reads corresponding to sequences not contained in the original library. Only reads corresponding the wild-type Acr and reads containing mutations in *exactly one* codon relative to wild-type were kept for further analysis. Read counts of the remaining reads were augmented with pseudocounts and normalized within each sorting fraction. Then, for each single mutant, read counts were normalized across fractions, resulting in a distribution of read counts over fractions for each mutant.

### Acr mutant activity regression

As a proxy for Acr activity, linear regression of the binned distribution and mean of log fluorescence intensities was performed for a set of single Acr mutants (benchmarks). To this end, log fluorescence intensity data from flow cytometry for the benchmark mutant set were binned (bin width 0.5, range from 0 to 12) and bins normalized to sum to one. An affine regression model relating the read distribution across sorting fractions to the distribution of log fluorescence intensity was zero initialized and fitted using gradient descent to minimize squared error under L^2^ regularization. Model fits were evaluated using mean square error calculated via leave-one-out cross-validation on the set of benchmark mutants.

Log fluorescence intensity distributions were predicted for all single Acr mutants. As a measure of confidence for the predicted distribution, the relative entropy of the underlying NGS read distribution was computed relative to a uniform distribution. Large relative entropy indicates a distribution with a majority of probability mass focused on one sequenced fraction (high confidence), whereas a low relative entropy indicates a distribution close to the uniform distribution (low confidence).

### Mammalian cell culture

HEK293T (human embryonic kidney) cells were cultured at 5% CO_2_ and 37°C in a humidified incubator and passaged every 2-3 days. Cells were maintained in DMEM (Thermo Fisher Scientific) supplemented with 10% (v/v) fetal calf serum (Thermo Fisher Scientific), 100 U mL^-1^ penicillin, 100 µg mL^-1^ streptomycin (Thermo Fisher Scientific). Prior to the assays, cells were checked for mycoplasma contamination by qPCR (Mycoplasma Check, GATC Eurofins).

12,500 cells were seeded in a 96 well culture plate and transfected the next day with Lipofectamine 3000 (Thermo Fisher Scientific) according to the manufacturer’s protocol. Cells were co-transfected with 33 ng of (i) a plasmid co-expressing *Renilla* and firefly luciferase as well as a sgRNA targeting the firefly reporter gene, (ii) 33 ng of a plasmid encoding *Spy*Cas9 with C- and N-terminal nuclear localization signals (NLS) and an N-terminal 3xFlag tag, (iii) and either 11 ng, 33 ng or 99 ng of a vector encoding human codon-optimized Acr mutant variants. A stuffer plasmid (pUC19) was used to top up plasmid levels per sample to 165 ng, thereby keeping the total amount of DNA constant across all samples. 72 hours post-transfection, cells were washed with PBS and firefly and *Renilla* luciferase activities were measured using the Dual-Glo luciferase assay system (Promega) according to manufacturer’s instructions. First, one volume Dual-Glo reagent was added to each well. Following a ten-minute incubation time, lysates were transferred to a white 96-well plate and firefly luciferase photon counts were measured using a FLUOstar Omega multimode reader (BMG Labtech). Subsequently, one volume Dual-Glo Stop & Glo was added to quench the firefly signal and activate *Renilla* luciferase, samples were incubated for ten minutes, and *Renilla* luciferase photon counts were measured. To calculate the reported luciferase activity values, firefly luciferase photon counts were normalized to *Renilla* luciferase photon counts in each sample.

### Protein production for Acr single point mutants

Bacterial codon-optimized sequences of AcrIIA4 mutants were PCR amplified and cloned into pET-28b(+) using BsaI sites. Constructs were expressed with an N-terminal His-tag. Plasmids were transformed into BL21(DE3) competent cells (Thermo Fisher Scientific). Protein expression was conducted in LB medium supplemented with kanamycin, and cells were induced with 0.5 mM IPTG at an OD_600_ of 0.6, followed by incubation at 18°C for 16 hours. Next, cells were harvested by centrifugation, resuspended in lysis buffer (50 mM Tris-HCL pH 7.5, 200 mM NaCl, 1 mM DTT, 1 mM PMSF, and 0.3 mg ml^-1^ lysozyme) and sonicated. Subsequently, samples were centrifuged, and the cleared lysates were incubated with 1 mL HisPur Ni-NTA Resin (Thermo Fisher Scientific), washed and eluted. Eluates were desalted using 7K MWCO Zeba Spin Desalting columns, stored in phosphate-buffered saline solution (PBS). The flow-through was collected and concentrated using 3K MWCO Pierce Protein Concentrators PES (Thermo Fisher Scientific), followed by flash freezing.

### Affinity measurements

Biolayer interferometry binding data were collected with a Gator Bioanalysis System and processed using the instrument’s integrated software. For the binding assay, His tagged designs were loaded onto anti-NTA biosensors (Gator probes) at 5 µg mL^-1^ in binding buffer (PBS pH 7.4) for 120 s. Homemade CRISPR-Cas9 RNP was diluted from 200 nM to 6.25 nM in binding buffer. After baseline measurement in the binding buffer alone, the binding kinetics were monitored by dipping the biosensors in wells containing the Cas9 RNP at the indicated concentration (association step) and then dipping the sensors back into baseline/buffer (dissociation).

## RESULTS

### Implementation and validation of an anti-CRISPR deep mutational scanning pipeline

Our anti-CRISPR deep mutational scanning pipeline comprises four steps: (1) generation of the Acr mutational library; (2) Implementation of a CRISPR interference (CRISPRi)-based gene circuit that can be inhibited by an Acr, hence enabling selection of Acrs mutants according to their activity; (3) FACS-mediated fractionation of the mutational library based on the intensity of a CRISPRi-dependent fluorescent reporter signal and deep amplicon sequencing of the individual fractions; (4) data analysis and integration using a mathematical model trained on a set of benchmark mutants (Figure 1, workflow).

**Figure 1:**
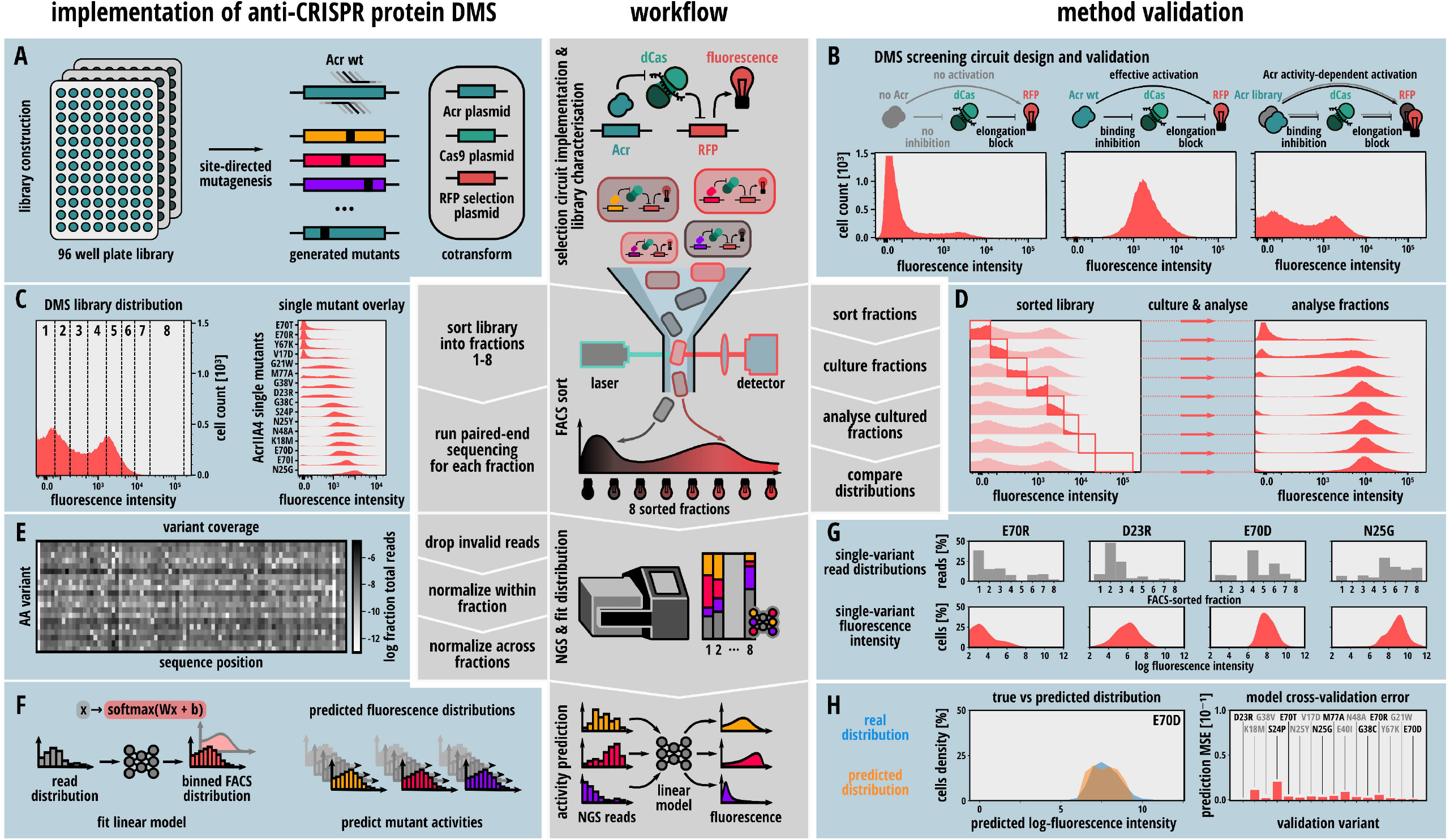
Implementing an anti-CRISPR protein deep mutational scanning pipeline. Center: General setup of the deep mutational scanning workflow comprising (i) a CRISPRi-based selection circuit in *E. coli* generating a fluorescent output indicating Acr activity, (ii) a Flow-seq pipeline consisting of enrichment of Acr mutants of distinct activity by FACS, followed by NGS, and (iii) a data analysis pipeline to predict inhibition potencies for every Acr mutant. (**A**) Schematic showing the generation of the Acr mutant library (left) and the three-plasmid system used for selection (right). (**B**) Validation of the CRISPRi gene circuit. *E. coli* carrying plasmids expressing an RFP reporter, d*Spy*Cas9, and an RFP gene-targeting sgRNA were transformed with a control vector (left), with a vector expressing wild-type AcrIIA4 (center) or with the AcrIIA4 mutant library (right), followed flow cytometry analysis. (**C**) Left: Eight fractions corresponding to the indicated bins were selected and 150,000 cells per fraction were sorted. Right: Flow cytometry measurement of the fluorescence distribution obtained for 16 AcrIIA4 mutants (indicated). Mutants were selected, so that they span the whole activity spectrum from very weak to very strong Cas9 inhibition. (**D**) Validation of successful sorting of AcrIIA4 variants with distinct inhibition properties corresponding to the eight fractions in **C**. For each fraction, cultures were re-grown overnight individually before flow cytometry analysis. (**E**) Analysis of AcrIIA4 mutant coverage. The heat map shows the proportion of reads for each Acr mutant from the total number of NGS reads. Reads from all fractions were pooled for this analysis. (**F**) A linear regression model was trained on the NGS read distribution profiles and corresponding fluorescence profiles of the 16 single mutants in **C**. The model was then applied to the NGS data of all other AcrIIA4 mutants to predict the underlying fluorescence profile and calculate the log mean fluorescence as a measure for Acr activity. (**G**) Exemplary NGS read distribution profiles and corresponding fluorescence profiles obtained by flow cytometry. (**H**) Left: Overlay of the model-predicted fluorescence distribution and the experimentally determined distribution exemplified for the E70D AcrIIA4 mutant. Right: Estimation of model accuracy by leave-one-out cross-validation using NGS and fluorescence profiles of the 16 individually tested AcrIIA4 mutants.

First, to implement a robust gene circuit for Acr selection, we created a three-plasmid system encoding (i) a (mutant) Acr protein, (ii) a catalytically impaired *Spy*Cas9 (d*Spy*Cas9) and sgRNA, and (iii) a red fluorescent protein (RFP) reporter (Figure 1A, right). The sgRNA was designed to direct the d*Spy*Cas9-sgRNA ribonucleoprotein complex to the 5’ region of the RFP reporter non-template strand, thereby efficiently blocking transcriptional elongation through CRISPRi (Figure 1B, left), as previously described (48). In presence of an Acr that prevents *Spy*Cas9 from binding its target DNA, RFP expression is released, as we determined by flow cytometry analysis (Figure 1B, center). While it is well-established that AcrIIA4 abolishes Cas9 DNA binding (19,49,50), the ability of AcrIIA5 to block DNA targeting by Cas9 has been under debate (20,21). When tested in context of our *E. coli* CRISPRi selection circuit, however, both AcrIIA4 (Figure 1B, center) and AcrIIA5 (Supplementary Figure S1A, wild-type AcrII5) showed a considerable rescue of reporter expression as compared to the CRISPRi control. This indicates that also AcrIIA5 is capable of considerably weakening Cas9 DNA binding, at least under our experimental conditions.

To create libraries covering all single point mutants for AcrIIA4 and -5, we performed PCRs with primer pairs that replace each individual codon with NNB randomized codons (Figure 1A). Due to the small size of Acrs typically around ∼100 amino acids, this straight-forward procedure is both rapid and cost-efficient. We first cloned the single codon sub-libraries into expression plasmids, then mixed the resulting plasmid sub-libraries at equimolar ratios, followed by transformation into *E. coli* DH5α receiver cells already carrying the two other CRISPRi selection plasmids (Figure 1A). From the transformed cells, cultures were grown to stationary phase, diluted and Acr and Cas9 expression was induced with arabinose and IPTG, respectively, followed by flow cytometry. The resulting *E. coli* AcrIIA4 and -5 libraries showed a broad distribution with respect to the RFP reporter fluorescence, indicating that they contained Acrs of various strengths (Figure 1B, right and Supplementary Figure S1A). Remarkably, the AcrIIA5 library showed a notable right-shift in the high-fluorescence peak as compared to wild-type AcrIIA5 (Supplementary Figure S1A), suggesting that the library contained some AcrIIA5 variants preventing Cas9 DNA binding more efficiently than the wild-type Acr.

We then performed flow sorting of the libraries and collected a total of eight fractions, which covered the entire fluorescence spectrum (Figure 1C, left and Supplementary Figure S1B). To determine whether sorted fractions indeed carried Acr variants of distinct strength, we re-grew cultures from each individual fraction separately and measured RFP expression by flow cytometry (Figure 1D and Supplementary Figure S1C). We observed distinct fluorescence distributions in particular for FACS fractions 1-5, which corresponded to the low to medium-high fluorescence fractions (Figure 1D and Supplementary Figure S1C). This suggested that each fraction was enriched for Acrs with distinct inhibition potency. We then sequenced the Acr mutant pools in all individual fractions on the Illumina MiSeq platform.

To analyze general mutant coverage, we first summed up all NGS reads in all fractions and calculated the proportion of reads corresponding to any given mutant (Figure 1E and Supplementary Figure S2). Overall, 93% of all possible AcrIIA4 and -5 mutants could be robustly detected within the library (>10 reads), and the majority of mutants were covered by more than 100 reads.

Next, to determine the inhibition potency for each Acr mutant in the library, we calculated the frequency of NGS reads corresponding to each Acr mutant in each fraction (Supplementary data 2 and 3). We then fitted an affine regression model to the obtained NGS read distributions (Figure 1F). The model comprised a fully parameterized affine transformation and was trained on NGS read frequencies and corresponding FACS fluorescence profiles of 16 individually measured benchmarks mutants for AcrIIA4 (Figure 1C, right) and 12 benchmark mutants for AcrIIA5 (Supplementary Figure S3, see Material & Methods for details). We then used the trained model to predict RFP fluorescence profiles corresponding to each Acr mutant within our *E. coli* library from the individual NGS read profiles (Figure 1G; see Supplementary data 4 and 5 for the predicted fluorescence profile for every single AcrIIA4 and -5 mutant, respectively). Leave-one-out cross-validation on the benchmark mutant dataset (Figure 1C, right) revealed that our model could accurately infer the reporter fluorescence distribution, and hence the activity of the Acr variant (Figure 1H). From the predicted fluorescence distribution, we finally calculated the log fluorescence intensity. This value corresponds to the model-predicted ability of the Acr variant to prevent CRISPRi as a measure for the Acr inhibition potency.

To cross-validate our analysis, we compared the obtained Acr inhibition potencies (log mean fluorescence values) with two previously reported, smaller AcrIIA4 mutant datasets (50,51) (Supplementary Figure S4). Despite the fact that the readouts underlying these studies were profoundly different from our DMS readout, namely measurement of *Spy*Cas9 gene drive activity in *Saccharomyces cerevisiae* in case of Basgall et al. (51) and *in vitro* DNA cleavage in case of the Dong et al. study (50), our DMS data correlated with both literature datasets reasonably well (R = -0.82 and 0.79 for the Basgall et al and Dong et al datasets, respectively; Supplementary Figure S4).

On top of predicting the Acr activity from the NGS profiles, we also estimated the confidence in each data point. To this end, we calculated the relative entropy, i.e. the distance of the NGS read count distribution over the different fractions from a uniform distribution, for each Acr mutant. Hence, for each mutant in our library we obtained two values, namely its predicted activity as well as the confidence in this prediction.

### Anti-CRISPR mutational fitness landscapes reveal mutational robustness

From our analysis, we obtained comprehensive mutational fitness landscapes for both, AcrIIA4 (Figure 2A) and AcrIIA5 (Supplementary Figure S5A).

**Figure 2:**
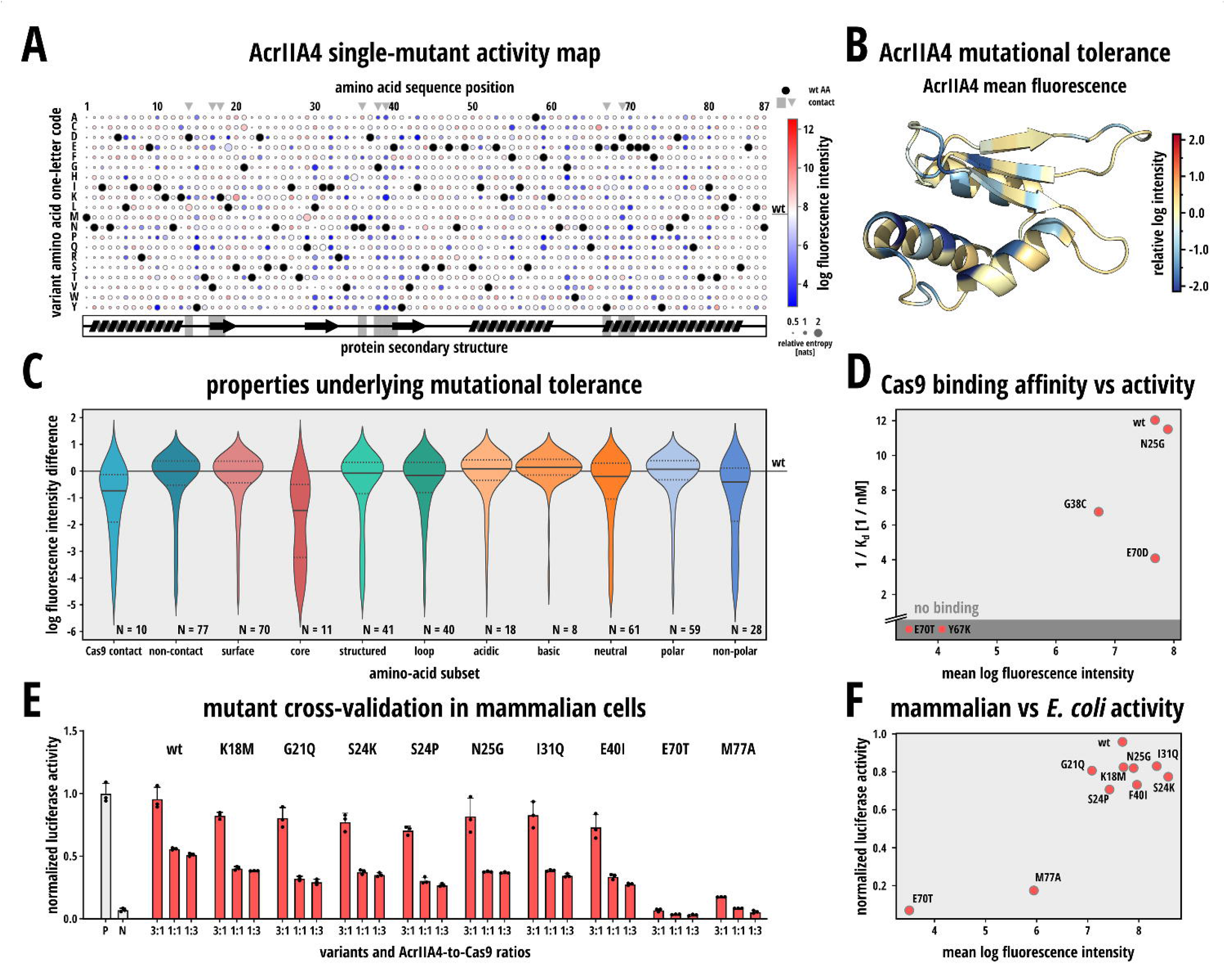
The AcrIIA4 mutational fitness landscape reveals high mutational tolerance. (**A**) AcrIIA4 mutational fitness landscape. Circle color indicates inhibition potency of the indicated mutant. Circle size corresponds to relative entsropy of the underlying NGS read distribution as a measure for the confidence. The relative entropy indicates the distance of the NGS read distribution from a uniform distribution (noise). Black circles correspond to the wild-type residue. Grey triangles and grey regions in the secondary structure cartoon below the heat map indicate residues directly interacting with Cas9. (**B**) For each residue the mutational tolerance, i.e. the mean activity across the 19 single mutants per position, was color-coded onto the AcrIIA4 structure (PDB 5VW1). (**C**) Violin plots showing the distribution of the predicted log mean fluorescence values for the indicated residue sub-groups. (**D**) Scatter plot of predicted log mean fluorescence values in **A** and corresponding Cas9 binding affinities measured by biolayer interferometry. (**E**) Cross-validation of inhibition potency for individual AcrIIA4 mutants in mammalian cells. HEK 293T cells were co-transfected with constructs expressing (i) *Renilla and* firefly luciferase as well as a sgRNA targeting the firefly reporter gene, (ii) *Spy*Cas9 and (iii) the indicated AcrIIA4 variant, followed by luciferase assay. The Cas9:Acr vector mass ratios used for transfection are indicated. (**F**) Scatter plot of predicted log mean fluorescence values in **A** and corresponding luciferase activities in **E** (for the Acr:Cas9 ratio of 3:1).

Very generally, AcrIIA4 and -5 both well-tolerate single point mutations at most positions throughout the protein sequence (Figure 2A,B and Supplementary Figure S5). More specifically, 79% and 74% of the mutated AcrIIA4 and -5 variants, respectively, showed an inhibition potency corresponding to at least 90% of their cognate wild-type Acr. In case of AcrIIA4, mutational tolerance was rather independent of whether mutations were introduced into loop regions or structured regions and also mostly independent of the chemical properties of the wild-type amino acid underlying the mutated residue (polar, acidic, basic) (Figure 2C). Mutations were generally less-well tolerated at the protein core as compared to the Acr surface (Figure 2C). However, Acr surface sites directly contacting Cas9 were rather mutation intolerant (Figure 2C), which is expected considering their key role in the Acr-Cas9 interaction. Still, also these residues usually tolerated various substitutions. The AcrIIA5 DMS data set was noisier as compared to the AcrIIA4 data as evident from the overall lower entropy (Supplementary Figure S6). For a subset of AcrIIA5 mutants in our library, however, we could still confidently measure their activity (Supplementary Figure S4A). Alike AcrIIA4, AcrIIA5 also well-tolerated substitutions at most residues throughout the protein (Supplementary Figure S5A and B). Notably, AcrIIA5 mutations in the protein core and the N-terminal intrinsically disordered region were tolerated less (Supplementary Figure S5B). This underlines the importance of the disordered protein region for AcrIIA5 inhibitory function. Moreover, the AcrIIA5 dataset contained various mutants that showed an activity higher than the wild-type Acr (Supplementary Figure S5A). This suggests that unlike AcrIIA4, which naturally prevents Cas9 DNA binding very effectively, the ability of AcrIIA5 to inhibit Cas9 DNA binding is naturally rather weak but might be enhanced by mutagenesis. Taken together, our DMS analysis of AcrIIA4 and -5 revealed that both CRISPR inhibitors show a considerable mutational robustness.

### Mutant activity determined by DMS corresponds with Cas9 binding affinity *in vitro* and cleavage inhibition in human cells

Next, to determine whether the mutation-induced modulation of Cas9 inhibition was related to changes in Cas9 binding affinity, we purified wild-type AcrIIA4 as well as five AcrIIA4 mutants of varying activity from our library (N25G, G38C, Y67K, E70D and E70T; Supplementary Figure S7). We then determined the binding affinity of these Acr variants to Cas9 RNP using biolayer interferometry (BLI) and compared the obtained affinities with the inhibition potencies derived from our DMS analysis (Figure 2D). While we were able to detect tight binding to Cas9 for high activity AcrIIA4 mutants, no binding was detected for Y67K and E70T, the two mutants with the lowest mean log fluorescence intensity in the affinity measurement test set (Figure 2D and Supplementary Figure S8). This indicated that the functional impact of mutation was mostly due to alterations in Cas9 binding affinity rather than other properties such as protein stability or expression levels.

Finally, to assess whether our *E. coli* DMS pipeline can inform the selection of Acrs with defined inhibition strength for applications in mammalian cells, we human codon optimized and expressed a subset of our AcrIIA4 mutants in HEK293T cells alongside a luciferase reporter, Cas9 and a luciferase-targeting sgRNA. We then compared the resulting luciferase activities (Figure 2E) with the Acr potencies (log mean fluorescence intensities) as measured by DMS in *E. coli* (Figure 2A). We found that AcrIIA4 mutants classified as active, moderately active or inactive in our DMS analysis also showed – at the qualitative level - corresponding activities in the mammalian cell assay (Figure 2F). This indicates that the mutational fitness landscapes resulting from our DMS analysis in *E. coli* can inform the selection of Acr mutants with desired properties for applications in other systems, including human cells.

## DISCUSSION

Here we present a powerful DMS pipeline to comprehensively sample the mutational fitness landscape of anti-CRISPR proteins and applied it to AcrIIA4 and -5, two prominent inhibitors of the widely-applied CRISPR-Cas9 from *Streptococcus pyogenes*. Very broadly, we found that both Acrs were highly tolerant against single-residue substitutions across most parts of the protein. This is in line with a recent, independent study by Figueroa and colleagues on an unrelated anti-CRISPR (AcrIF7), for which 2/3 of the *in vitro* tested single mutations were well-tolerated, despite considerable evolutionary conservation of several of these residues (27).

Interestingly, we observed that even residues that lie within known, mechanistically important Acr regions, e.g. the Cas9-contacting loops in AcrIIA4 (49,50) or the intrinsically disordered N-terminus of AcrIIA5 (52), still tolerate mutations, at least to some extent. This suggests that a certain degree of “intrinsic redundancy” with respect to the Acr-Cas effector contacts that form the basis of the Cas inhibitory activity of AcrIIA4 and -5.

On the one hand, these findings are exciting from an evolutionary standpoint. The mutational robustness might be a consequence of selection pressure to render the Acr functional despite high mutation frequencies in phages. Mutational tolerance might also be important for Acr plasticity and hence enable adaptation of Acrs to alterations of the cognate Cas effector structure. It might also aid Acr adaption to changes in cellular context or external conditions that impact Acr folding (e.g. pH, temperature).

On the other hand, these observations are also relevant from a protein engineering and CRISPR application standpoint. Since many Acrs robustly function across a wide range of cell types, they present interesting tools for controlling, fine-tuning and safeguarding the activity of CRISPR-Cas effectors used in biotechnology or medicine (53). Acrs have, for instance, been used to limit the time-window of Cas9 activity or to fine-tune Cas9 activity in human cells, thereby reducing off-target editing (30,31). Moreover, miRNA-dependent Acr transgenes have recently been expressed in cells (35) and mice (36), thereby restricting CRISPR-Cas9-meditated genome editing to selected cell types. Finally, the ability to effectively shut off CRISPR-Cas effectors could be important in context of ecosystem engineering with gene drives (51,54). For such exciting applications, strategies to enhance or attenuate the inhibition potency of Acrs by mutagenesis will likely be of future interest. Moreover, for their *in vivo* application, the ability to alter potentially immunogenic epitopes in Acrs without perturbing their function, could also become relevant. Finally, mutational fitness landscapes obtained by DMS will facilitate the optimization of Acrs with respect to their inhibition potency, as we exemplified for AcrIIA5 through the identification of candidate mutations that resulted in a more potent Cas9 inhibition under our experimental conditions.

While a number of previous studies characterized mutant Acrs, including variants of AcrIIA4 (49-51) and AcrIIA5 (52), these analyses were either focused on a few, selected residues or limited types of substitution or truncation. In contrast, our Acr-DMS pipeline provides a detailed and differentiated picture of the Acr mutational fitness landscapes. Our data shows that regions in AcrIIA4 and-5 considered important for CRISPR-Cas inhibition tolerate at least a subset of possible substitutions, provided the remainder of the Acr remains intact. In the future, it will be possible to extend our DMS pipeline to other CRISPR-Cas orthologues and Acr types and families. While we employed a gene circuit based on CRISPRi as readout, since it could be easily implemented in case of *Spy*Cas9 and provided a decent dynamic range of measurement under our experimental conditions, it should also be possible to set up selection circuits based on DNA or RNA target cleavage or even Cas effector binding. Taken together, our study establishes Acr mutational fitness landscapes created via deep mutational scanning as powerful resource to study the Acr evolutionary design space and inform the engineering of Acrs that are optimized for selected applications in CRISPR genome engineering.

## DATA AVAILABILITY

Plasmid sequences (Genbank files) are provided as Supplementary data 1. NGS read distributions for AcrIIA4 and -5 are available as Supplementary data 2 and 3, respectively. Predicted flow cytometry plots for each single AcrIIA4 and -5 mutant are available as Supplementary data 4 and 5, respectively. Code is available on GitHub (https://github.com/mjendrusch/acr-dms) and the entire datasets are accessible on Zenodo (https://dx.doi.org/10.5281/zenodo.5221560). Data will also be made available from the corresponding author upon reasonable request.

## ACKNOWLEDGEMENTS

We thank members of the Niopek and Correia labs at TU Darmstadt and EPFL Lausanne, as well as Max Waldhauer and Stefan Holderbach, Heidelberg University, for helpful discussions. We would like to thank the Protein Production and Structure Core facility at EPFL for their support on the protein biophysical characterization experiments, the ZMBH flow cytometry core facility (Heidelberg University) for help with bacterial cell sorting and the EMBL Genomics Core Facility (EMBL Heidelberg) for performing amplicon deep sequencing.

## Author contributions

D.H. and D.N. conceived the initial idea and refined it together with T.S. and M.J.. T.S. and D.H. designed and performed experiments and purified the Acrs. S.R. performed affinity measurements. M.J. conceived and implemented the mathematical model for analysis of FACS/NGS data and performed correlation analysis. B.E.C. provided critical expertise on affinity measurements and data interpretation. D.N. directed the work and secured funding. T.S., D.H., M.J. and D.N. wrote the manuscript with support from all authors.

