## Supplementary Figures and Tables for "A deep mutational scanning platform to characterize the fitness landscape of anti-CRISPR proteins"

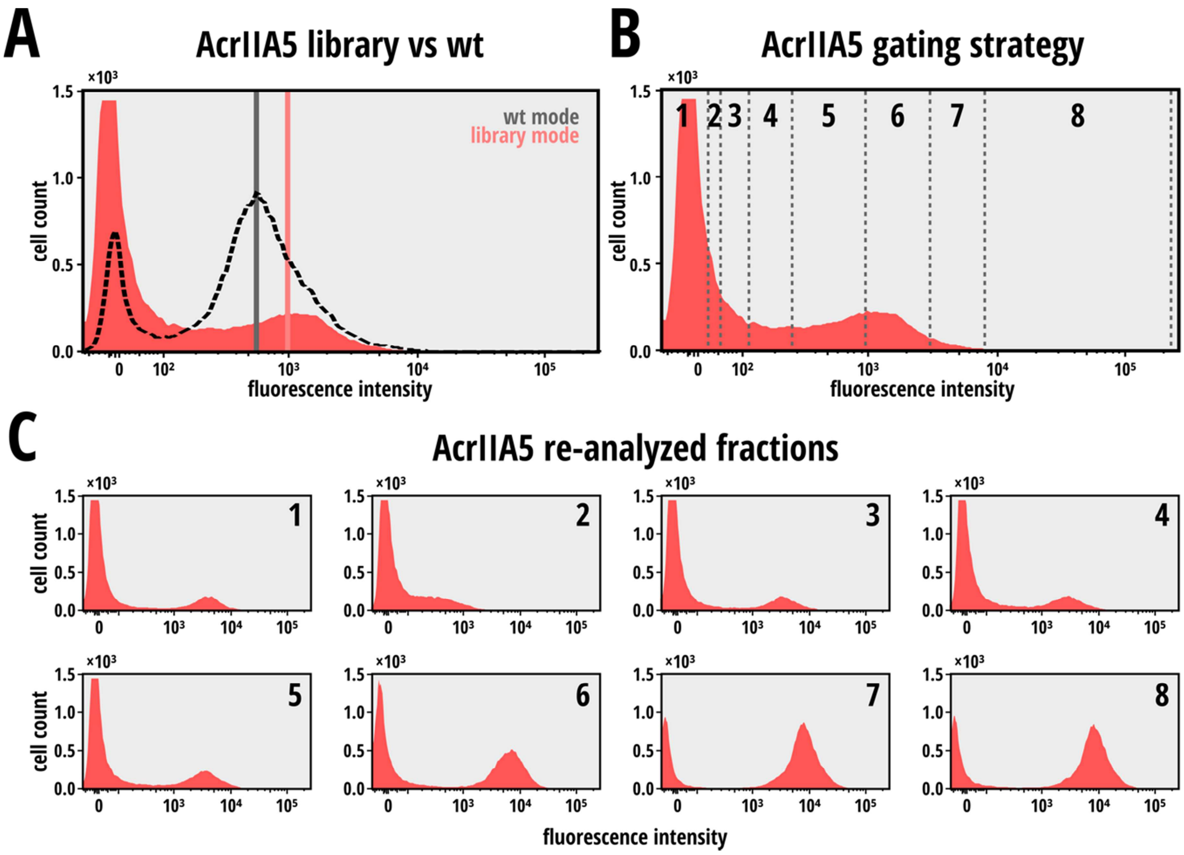


**Supplementary Figure S1: FACS sorting of the AcrIIA5 mutation library.** (**A**) Overlay histogram showing the RFP profile of the AcrIIA5 mutant library in comparison to that of wild-type AcrIIA5. (**B**) Eight fractions corresponding to the indicated bins were selected and 150,000 cells per fraction were sorted. (**C**) Validation of successful sorting of AcrIIA5 variants with distinct inhibition properties corresponding to the eight fractions in **B**. For each fraction, cultures were re-grown overnight individually before flow cytometry analysis.

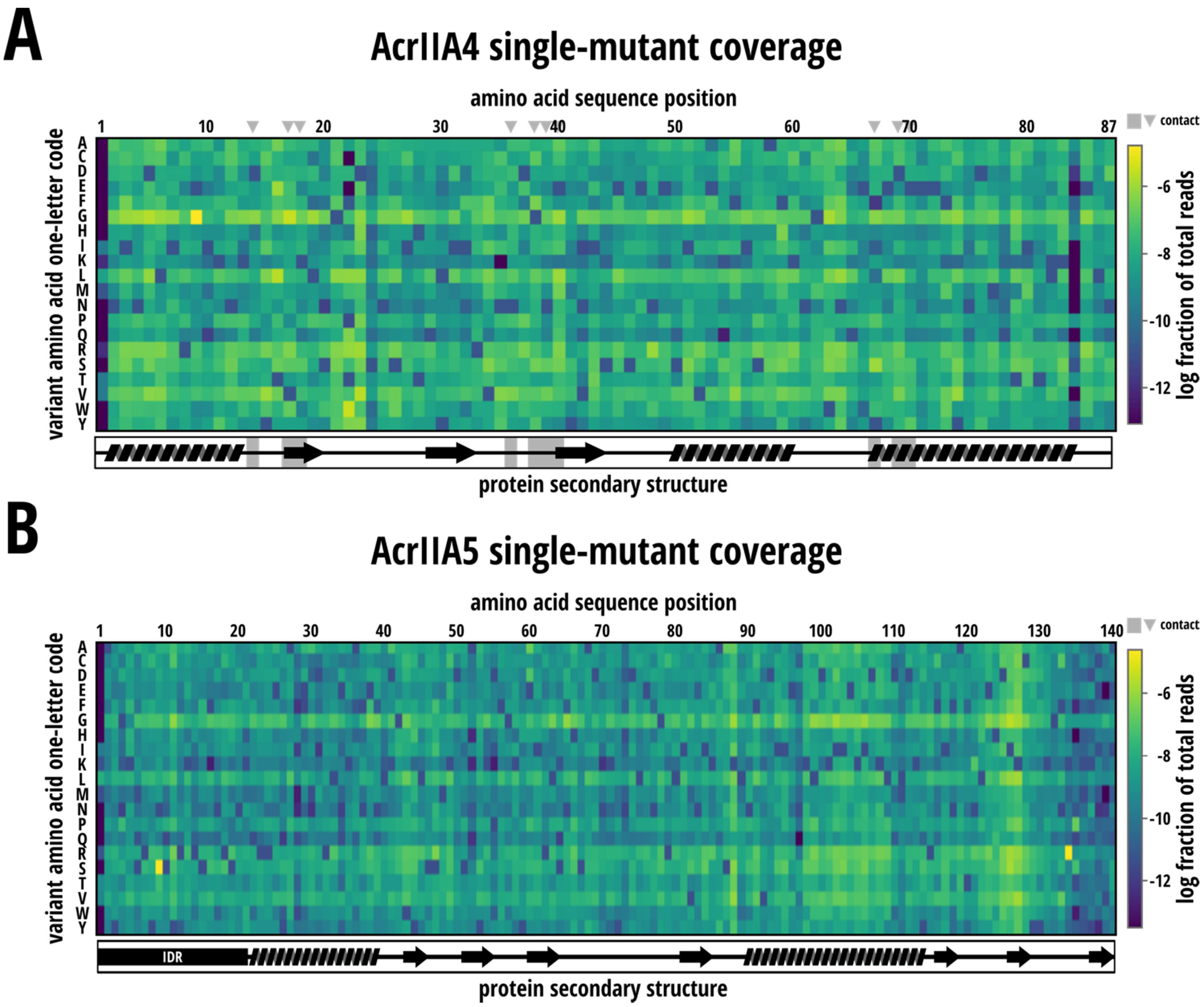


**Supplementary Figure S2: Library coverage.** Reads from all fractions were summed up for each single AcrIIA4 (A) or AcrIIA5 (B) mutant. The heat map indicates the log fraction of these sums from all reads for each mutant.

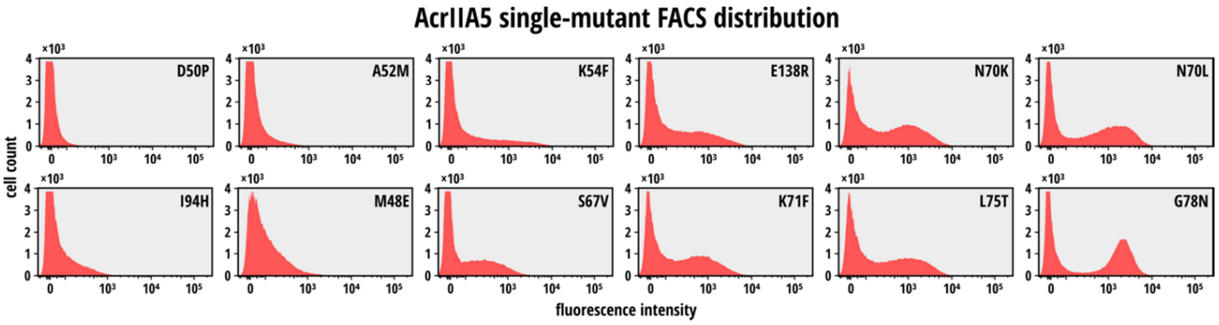


**Supplementary Figure S3: AcrIIA5 benchmark mutant fluorescence profiles.**
Fluorescence profiles corresponding to 12 individual AcrIIA5 mutants used for model training were obtained by flow cytometry. Mutants were selected, so that they span the whole activity spectrum from very weak to very strong Cas9 inhibition.

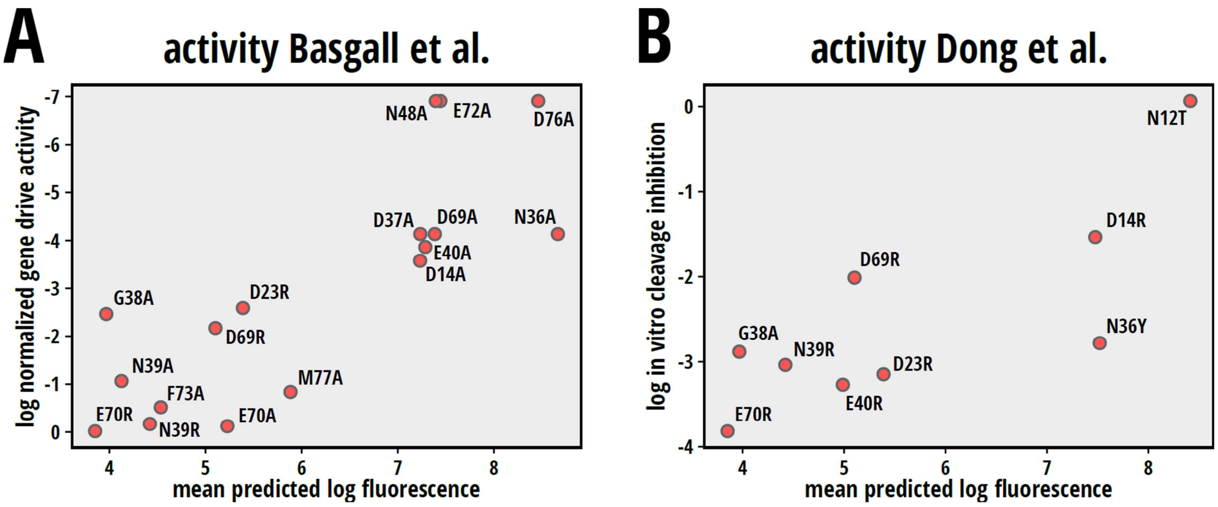


**Supplementary Figure S4: Cross-validation of DMS data sets with previously reported measurements of Acr mutant activity.** Scatter plots of the predicted mean log fluorescence derived from our DMS analysis and (**A**) the corresponding gene drive inhibition in *S. cerevisiae* (Basgall et al, 2018) or (**B**) the relative *in-vitro* DNA cleavage inhibition (Dong et al, 2017) for the indicated mutants.

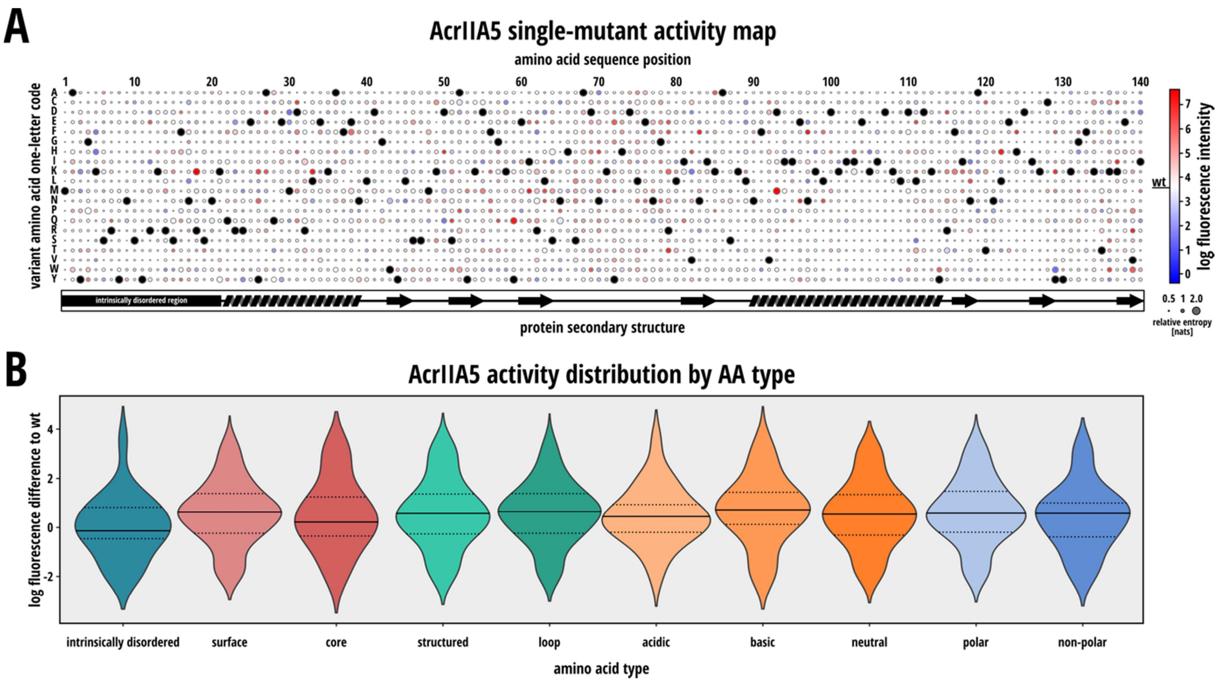


**Supplementary Figure S5: AcrIIA5 mutational fitness landscape.** (**A**) Circle color indicates inhibition potency (model-predicted log fluorescence intensity) for each AcrIIA5 mutant. Circle size indicates the entropy as a measure for the confidence. The entropy estimates the distance of the NGS read distribution form a uniform distribution (noise). Black circles correspond to the wild-type residue. The cartoon below the heat map indicates secondary structures. (**B**) Violin plots showing the distribution of the predicted log mean fluorescence values for the indicated residue sub-groups.

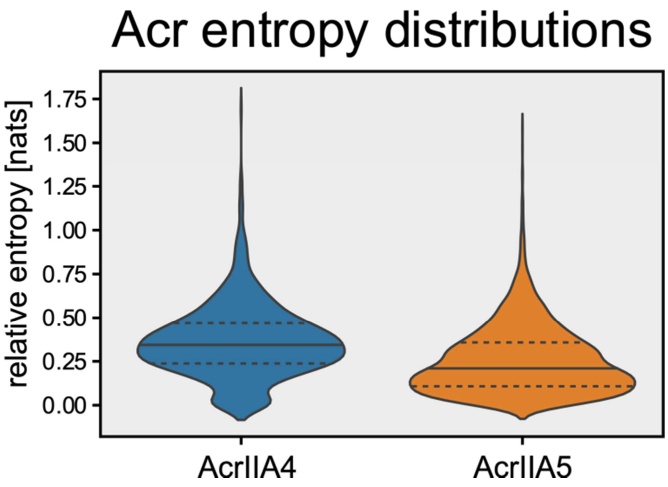


**Supplementary Figure S6**: Violin plots of the entropy distributions corresponding to the AcrIIA4 and AcrIIA5 DMS data sets.

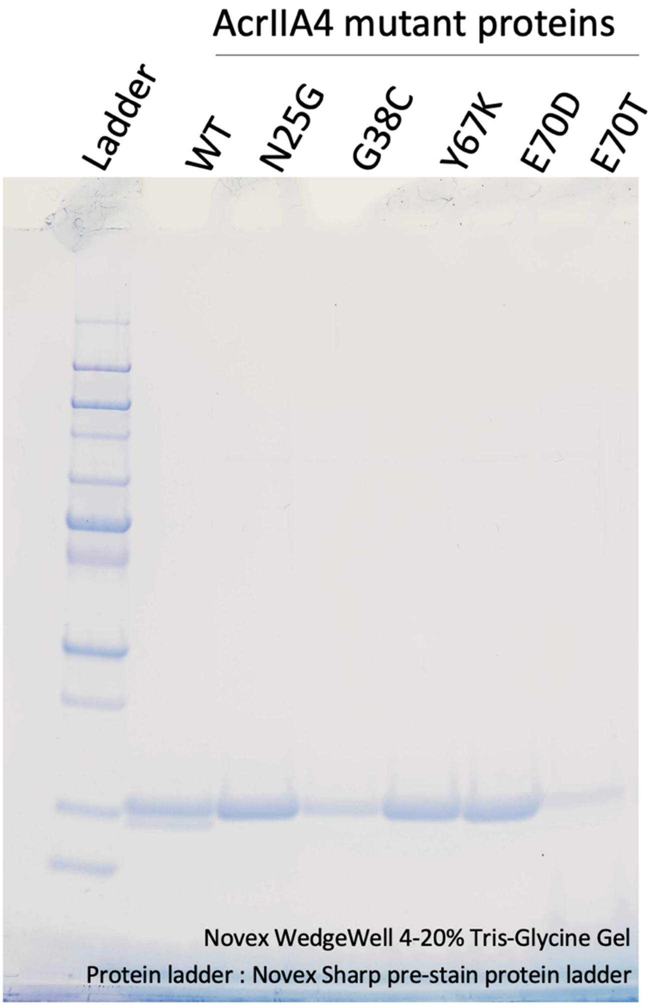


**Supplementary Figure S7:** Coomassie-stained SDS page gel of purified AcrIIA4 mutants.

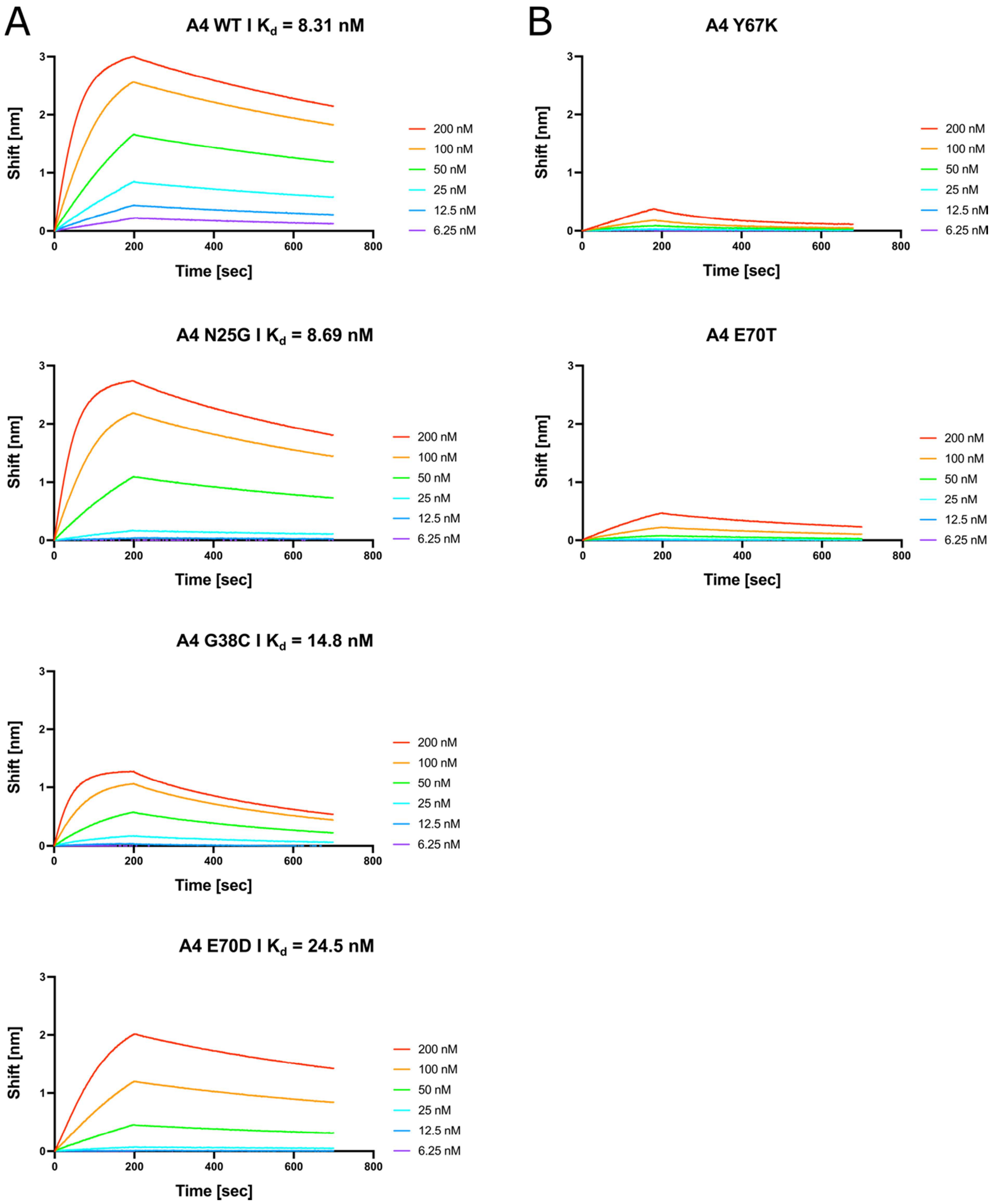


**Supplementary Figure S8:** Measurements of Cas9 binding affinity for AcrIIA4 variants. Biolayer Interferometry binding data from Gator probes at the indicated, different Acr proteins concentrations. While strong Cas9 RNP binding could be determined for the mutants in **A**, the mutants in **B** did not bind Cas9.

**Supplementary Table 1: List of plasmids used in this study.**

Amp = Ampicillin resistance, Kan = Kanamycin resistance, Cm = Chloramphenicol resistance, Ori = Origin of replication, ColE1 = high copy origin of replication (high copy), pSC101 = low copy origin of replication, p15A = low copy origin of replication, sfGFP = superfolder GFP, M0052 = M0052 degradation tag, CMV = cytomegalovirus, BGH = bovine growth hormone

**Plasmids for bacterial gene circuits:**

| **PlasmidID** | **Name** | **Resistance** | **Ori** | **Description** |
|---|---|---|---|---|
| 1 | pJUMP27-1A(sfGFP) | Kan | pSC101 | Addgene #126974, backbone for RFP expression plasmid |
| 2 | pBbA5c-RFP | Cm | P15A | Addgene #35281, backbone for d*Spy*Cas9 expression plasmid |
| 3 | pBAD24-sfGFPx1 | Amp | ColE1 | Addgene #51558, backbone for Acr expression plasmid |
| 4 | pBAD24_AcrIIA4 | Amp | ColE1 | Arabinose inducible expression of AcrIIA4, template plasmid for AcrIIA4 library generation |
| 5 | pBAD24_AcrIIA5(phageD4276) | Amp | ColE1 | Arabinose inducible expression of AcrIIA5, template plasmid for AcrIIA4 library generation |
| 6 | pJUMP27_J23102_RFP_M0052 | Kan | pSC101 | Constitutive RFP reporter expression with M0052 degradation tag |
| 7 | pBbA5C_sgMCS | Cm | P15A | Empty vector with additional MCS for sgRNA cassette |
| 8 | pBbA5C_sgMCS_dSpyCas9 | Cm | P15A | IPTG inducible expression of d*Spy*Cas9, additional MCS for sgRNA cassette |
| 9 | pBbA5C_sgMCS_dSpyCas9_RFPguide | Cm | P15A | IPTG inducible expression of d*Spy*Cas9, sgRNA cassette with spacer targeting RFP |
| 10 | pBAD24_AcrIIA4_V17D | Amp | ColE1 | AcrIIA4 single mutant, used for model training |
| 11 | pBAD24_AcrIIA4_K18M | Amp | ColE1 | AcrIIA4 single mutant, used for model training |
| 12 | pBAD24_AcrIIA4_G21W | Amp | ColE1 | AcrIIA4 single mutant, used for model training |
| 13 | pBAD24_AcrIIA4_D23R | Amp | ColE1 | AcrIIA4 single mutant, used for model training |

Supplementary Table 1 continued

| | | | | |
|---|---|---|---|---|
| 14 | pBAD24_AcrIIA4_S24P | Amp | ColE1 | AcrIIA4 single mutant, used for model training |
| 15 | pBAD24_AcrIIA4_N25G | Amp | ColE1 | AcrIIA4 single mutant, used for model training |
| 16 | pBAD24_AcrIIA4_N25Y | Amp | ColE1 | AcrIIA4 single mutant, used for model training |
| 17 | pBAD24_AcrIIA4_G38C | Amp | ColE1 | AcrIIA4 single mutant, used for model training |
| 18 | pBAD24_AcrIIA4_G38V | Amp | ColE1 | AcrIIA4 single mutant, used for model training |
| 19 | pBAD24_AcrIIA4_E40I | Amp | ColE1 | AcrIIA4 single mutant, used for model training |
| 20 | pBAD24_AcrIIA4_N48A | Amp | ColE1 | AcrIIA4 single mutant, used for model training |
| 21 | pBAD24_AcrIIA4_Y67K | Amp | ColE1 | AcrIIA4 single mutant, used for model training |
| 22 | pBAD24_AcrIIA4_E70D | Amp | ColE1 | AcrIIA4 single mutant, used for model training |
| 23 | pBAD24_AcrIIA4_E70R | Amp | ColE1 | AcrIIA4 single mutant, used for model training |
| 24 | pBAD24_AcrIIA4_E70T | Amp | ColE1 | AcrIIA4 single mutant, used for model training |
| 25 | pBAD24_AcrIIA4_M77A | Amp | ColE1 | AcrIIA4 single mutant, used for model training |
| 26 | pBAD24_AcrIIA5_G4P | Amp | ColE1 | AcrIIA5 single mutant, used for model training |
| 27 | pBAD24_AcrIIA5_S6Q | Amp | ColE1 | AcrIIA5 single mutant, used for model training |
| 29 | pBAD24_AcrIIA5_Q28Y | Amp | ColE1 | AcrIIA5 single mutant, used for model training |
| 30 | pBAD24_AcrIIA5_M48E | Amp | ColE1 | AcrIIA5 single mutant, used for model training |
| 31 | pBAD24_AcrIIA5_D50P | Amp | ColE1 | AcrIIA5 single mutant, used for model training |
| 32 | pBAD24_AcrIIA5_S51W | Amp | ColE1 | AcrIIA5 single mutant, used for model training |
| 33 | pBAD24_AcrIIA5_A52M | Amp | ColE1 | AcrIIA5 single mutant, used for model training |
| 34 | pBAD24_AcrIIA5_K54F | Amp | ColE1 | AcrIIA5 single mutant, used for model training |
| 35 | pBAD24_AcrIIA5_S67V | Amp | ColE1 | AcrIIA5 single mutant, used for model training |
| 36 | pBAD24_AcrIIA5_N70K | Amp | ColE1 | AcrIIA5 single mutant, used for model training |
| 37 | pBAD24_AcrIIA5_N70L | Amp | ColE1 | AcrIIA5 single mutant, used for model training |

Supplementary Table 1 continued

| | | | | |
|---|---|---|---|---|
| 38 | pBAD24_AcrIIA5_K71F | Amp | ColE1 | AcrIIA5 single mutant, used for model training |
| 39 | pBAD24_AcrIIA5_L75T | Amp | ColE1 | AcrIIA5 single mutant, used for model training |
| 40 | pBAD24_AcrIIA5_G78N | Amp | ColE1 | AcrIIA5 single mutant, used for model training |
| 41 | pBAD24_AcrIIA5_I94H | Amp | ColE1 | AcrIIA5 single mutant, used for model training |
| 42 | pBAD24_AcrIIA5_E138R | Amp | ColE1 | AcrIIA5 single mutant, used for model training |

**Plasmids for cell culture assays**

| **Plasmid ID** | **Name** | **Description** |
|---|---|---|
| 46 | ssAAV-SV40 hRluc-TK hLuc- H1-gRNA hLuc | SV40 promoter, *Renilla* luciferase; TK promoter, firefly luciferase, H1 promoter sgRNA targeting firefly luciferase |
| 47 | 3xFlag-NLS-SpyCas9-NLS | CMV promoter, 3xFlag-NLS-SpyCas9-NLS, BGH polyA |
| 48 | pCMV_AcrIIA4 | CMV promoter, AcrIIA4 wildtype, BGH polyA |
| 49 | pCMV_AcrIIA4_K18M | CMV promoter, AcrIIA4 mutant K18M, BGH polyA |
| 50 | pCMV_AcrIIA4_G21Q | CMV promoter, AcrIIA4 mutant G21Q, BGH polyA |
| 51 | pCMV_AcrIIA4_S24K | CMV promoter, AcrIIA4 mutant S24K, BGH polyA |
| 52 | pCMV_AcrIIA4_S24P | CMV promoter, AcrIIA4 mutant S24P, BGH polyA |
| 53 | pCMV_AcrIIA4_N25G | CMV promoter, AcrIIA4 mutant N25G, BGH polyA |
| 54 | pCMV_AcrIIA4_I31Q | CMV promoter, AcrIIA4 mutant I31Q, BGH polyA |
| 55 | pCMV_AcrIIA4_E40I | CMV promoter, AcrIIA4 mutant E40I, BGH polyA |
| 56 | pCMV_AcrIIA4_E70T | CMV promoter, AcrIIA4 mutant E70T, BGH polyA |
| 57 | pCMV_AcrIIA4_M77A | CMV promoter, AcrIIA4 mutant M77A, BGH polyA |

**Plasmids for protein purification**

| **Plasmid ID** | **Name** | **Description** |
|---|---|---|
| 58 | pET28a_AcrIIA4_wt | T7 Promoter, AcrIIA4 wildtype, T7 Terminator |
| 59 | pET28a_AcrIIA4_E70D | T7 Promoter, AcrIIA4 E70D mutant, T7 Terminator |
| 60 | pET28a_AcrIIA4_E70T | T7 Promoter, AcrIIA4 E70T mutant, T7 Terminator |
| 61 | pET28a_AcrIIA4_N25G | T7 Promoter, AcrIIA4 N25G mutant, T7 Terminator |
| 62 | pET28a_AcrIIA4_G38C | T7 Promoter, AcrIIA4 G38C mutant, T7 Terminator |
| 63 | pET28a_AcrIIA4_Y67K | T7 Promoter, AcrIIA4 Y67K mutant, T7 Terminator |

**Supplementary Table 2**: sgRNA target sites. Sequences in 5’-3’ direction, PAM sequences in bold.

| sgRNA target sites | Description |
|---|---|
| AACTTTCAGTTTAGCGGTCT**GGG** | RFP targeting sgRNA for CRISPRi gene circuit in *E. coli* cells |
| GGTAGTCGGTCTTAGAGTCC**AGG** | hLuc targeting sgRNA for cell culture validation |

**Supplementary Table 3: AcrIIA4 and -5 libraries overview**

| Library ID | Name | Description | Complexity (N x library) |
|---|---|---|---|
| Library 1 | pBAD24_AcrIIA4_mutant_library | Arabinose inducible expression of AcrIIA4 mutant library | 235 |
| Library 2 | pBAD24_AcrIIA5_mutant_library | Arabinose inducible expression of AcrIIA5 mutant library | 359 |

**Supplementary Table 4: Bacterial strains generated in this study**

| Strain ID | **Plasmid 1** Resistance = Amp Ori = ColE1 Arabinose inducible Acr expression | **Plasmid 2** Resistance = Kan Ori = pSC101 Lac-inducible d*Spy*Cas9 expression | **Plasmid 3** Resistance = Cm Ori = P15A Constitutive RFP expression | **Description** |
|---|---|---|---|---|
| I | 1 - pJUMP27-1A(sfGFP) | 8 - pBbA5C_sgMCS_dSpyCas9_RFPguide | 6 - pJUMP27_J23102_RFP_M0052 | Active CRISPRi system, no Acr expression (control) |
| II | 4 - pBAD24_AcrIIA4 | 8 - pBbA5C_sgMCS dSpyCas9_RFPguide | 6 - pJUMP27_J23102_RFP_M0052 | Active CRISPRi system, suppressed by wt AcrIIA4 expression |
| III | 5 - pBAD24_AcrIIA5 | 8 - pBbA5C_sgMCS dSpyCas9_RFPguide | 6 - pJUMP27_J23102_RFP_M0052 | Active CRISPRi system, suppressed by wt AcrIIA5 expression |
| IV | Library 1 – pBAD24_AcrIIA4_mutant_library | 8 - pBbA5C_sgMCS dSpyCas9_RFPguide | 6 - pJUMP27_J23102_RFP_M0052 | Active CRISPRi system, partially suppressed by AcrIIA4 library expression |
| V | Library 2 – pBAD24_AcrIIA5_mutant_library | 8 - pBbA5C_sgMCS dSpyCas9_RFPguide | 6 - pJUMP27_J23102_RFP_M0052 | Active CRISPRi system, partially suppressed by AcrIIA5 library expression |
