## Supplementary figures and images for "A deep mutational scanning platform to characterize the fitness landscape of anti-CRISPR proteins"

### Supplementary data 2

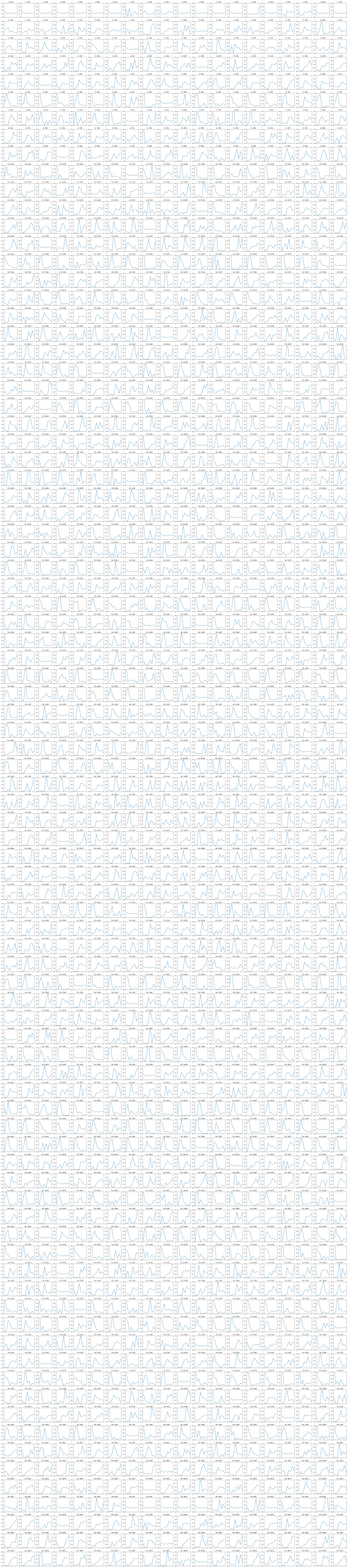

### Supplementary data 3

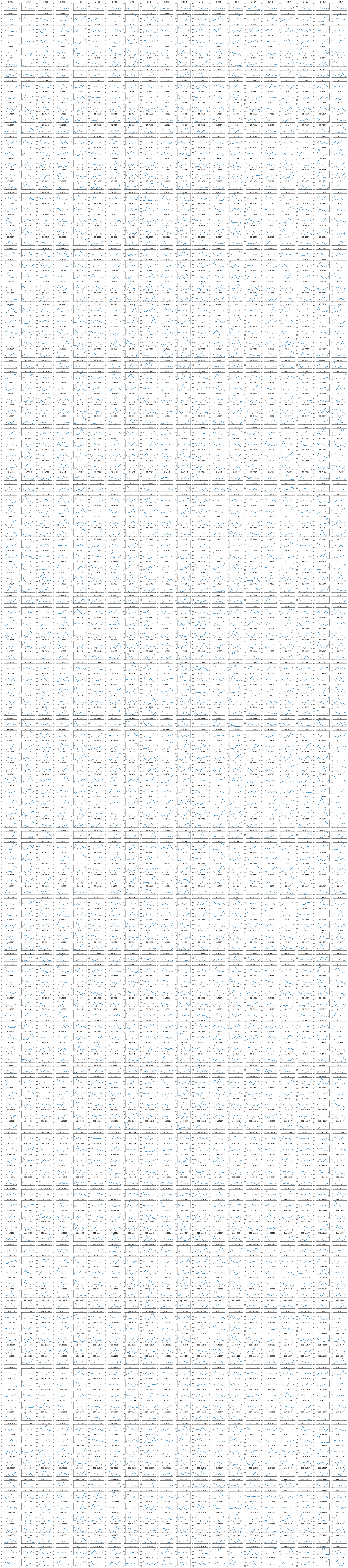

### Supplementary data 4

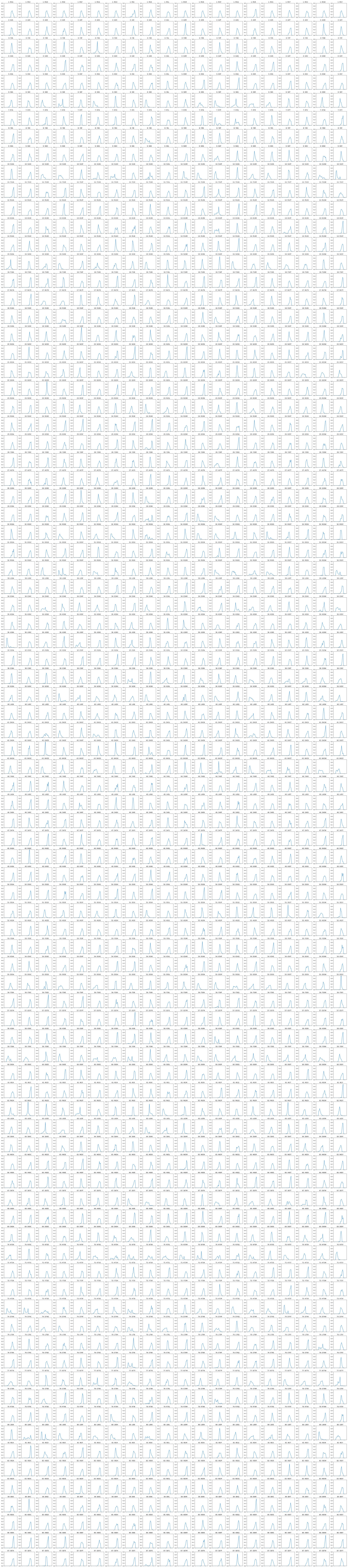

### Supplementary data 5

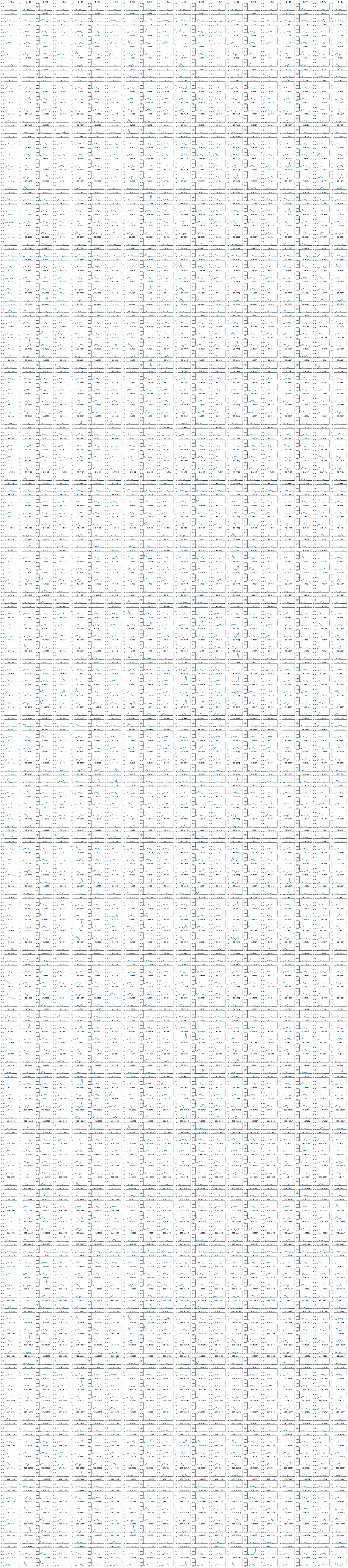
